# Blind De Novo Design of Dual Cyclic Peptide Agonists Targeting GCGR and GLP1R

**DOI:** 10.1101/2025.06.06.658268

**Authors:** Qiuzhen Li, Elisee Wiita, Thomas Helleday, Patrick Bryant

## Abstract

Agonists of the glucagon receptor (GCGR) and glucagon-like peptide-1 receptor (GLP1R) are key to treating metabolic diseases like type 2 diabetes and obesity, but often limited by being linear. Cyclic peptides offer greater stability and potential oral delivery, but their de novo design with computational methods as agonists is unproven. Here, we used the EvoBind AI platform to design cyclic peptide agonists for GCGR from sequence alone, without any structural templates or prior binding site information. EvoBind consistently identified the receptor’s activation surface, producing peptides that also activated GLP1R. Of three synthesised peptides, two showed potent dual activation of GCGR and GLP1R in cAMP assays, while all three activate GLP1R, with the 18-residue design achieving EC_50_ values of 32 nM (GLP1R) and 542 nM (GCGR), rivalling natural hormones. The designed sequences show no similarity to known agonists, and structural modelling indicates that they adopt novel binding modes compatible with active-state stabilisation. This is made possible by EvoBind’s purely sequence-based, template-free approach, which enables the discovery of alternative solutions that are inaccessible to structure-guided methods. This study presents the first successful blind *de novo* design of a cyclic peptide agonist for a G protein-coupled receptor (GPCR), using only the receptor sequence as input. The findings highlight EvoBind’s capacity to go beyond static binding, offering a generalisable strategy for therapeutic discovery through target-agnostic, sequence-driven peptide design.

## Introduction

Agonists targeting the glucagon-like peptide-1 receptor (GLP1R) and the glucagon receptor (GCGR) have transformed the treatment landscape for metabolic disorders, including type 2 diabetes and obesity. GLP1R, a 53 kDa class B G protein-coupled receptor (GPCR), is naturally activated by the 31-amino-acid incretin hormone GLP1, which is secreted in response to food intake and promotes glucose-dependent insulin secretion [1–3]. GCGR, a closely related receptor, plays a complementary role in glucose metabolism by mediating hepatic glucose output in response to glucagon. Therapeutic agents that act on both GLP1R and GCGR have shown promise for achieving enhanced metabolic control through synergistic effects on energy balance, glycaemic regulation, and weight loss [4].

A major limitation of current peptide-based therapies is their requirement for injection and limited oral bioavailability due to degradation by proteolytic enzymes, including dipeptidyl peptidase-4 [5,6]. Cyclisation of peptides offers a potential solution by increasing proteolytic stability and plasma half-life, which may support oral delivery [6,7]. However, designing cyclic peptides that retain agonist activity is particularly challenging for class B GPCRs, whose ligand-binding sites typically accommodate extended linear peptide conformations [8]. In the case of GLP1R and GCGR, a successful agonist must (1) bind with sufficient affinity, (2) fit within the confined activation pocket, and (3) induce receptor activation through conformational rearrangement of the transmembrane domain.

To date, most efforts have focused on engineering linear GLP1 analogues or hybrid sequences to improve potency or pharmacokinetics [8]. Small-molecule agonists have also been pursued but have not yet reached clinical approval [9]. Recent advances in computational protein design enable the generation of bespoke binders to protein targets [10–14], though these methods typically require detailed structural knowledge, such as holo-state models or annotated binding sites. EvoBind [15] represents a notable exception: it generates cyclic peptide binders using only the sequence of the target protein. While EvoBind has previously been shown to produce high-affinity cyclic binders and antagonists [16] in a structure-free manner, it remains untested whether such methods can also generate agonists, especially for GPCRs, without any structural or ligand-binding information.

Based on the rationale that AI models for structure prediction have learned suitable binding regions and that these are likely to be functional [17], we hypothesise that agonists can be designed without providing a design framework with a target region. Here, we explore this question by designing cyclic peptide agonists targeting GCGR using only its amino acid sequence as input to EvoBind. We proceed with testing the resulting designs for agonist activity on both GCGR and GLP1R, aiming to assess functional cross-reactivity and the potential for dual-acting therapeutics. Functional validation was performed using cAMP accumulation assays in cell lines expressing either receptor, and β-arrestin recruitment assays for GLP1R. This work represents the first demonstration of blind cyclic peptide agonist design for class B GPCRs and highlights the potential of sequence-based generative frameworks to identify and exploit functional receptor surfaces without structural guidance.

## Results

### Blind design of cyclic peptide GPCR agonists

Blind agonist design for a GPCR requires identifying an activation-competent surface and generating a ligand capable of inducing receptor activation, without structural templates or ligand information. Here, we additionally constrained the design space to cyclic peptides to improve the potential proteolytic stability and oral bioavailability. EvoBind [15], a design framework based on AlphaFold2 (AF2)[18] and AlphaFold-multimer (AFM)[19], was provided only with the amino acid sequence of the human glucagon receptor (GCGR; PDB 8JIQ) and tasked with generating de novo cyclic agonists (**Figure 1a**).

**Figure 1.**
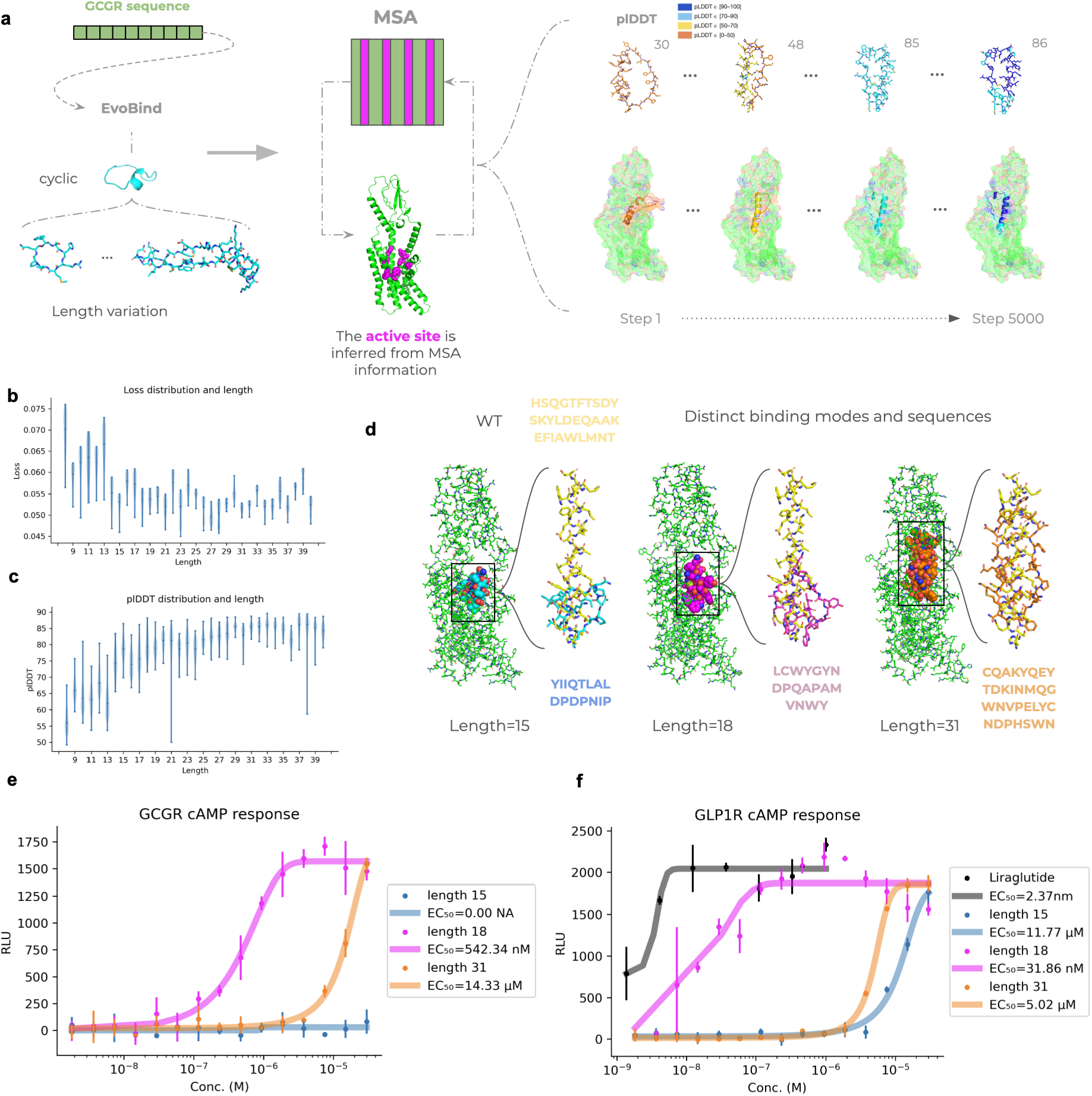
De novo design of cyclic peptide GPCR agonists using EvoBind. **a)** We used the design framework EvoBind to generate cyclic peptide agonists targeting GCGR in a fully automated, template-free manner. The design protocol runs for 5000 mutation steps per peptide, iteratively refining both sequence and structure. EvoBind identifies potential activation interfaces directly from the multiple sequence alignment (MSA) of the receptor, without prior knowledge of ligand binding sites. From the MSA, residues with strong co-evolutionary signals are highlighted as likely contact points, guiding the placement and orientation of the peptide. Throughout the design process, the model optimises interactions between the peptide and receptor, with the binding interface emerging naturally from evolutionary and structural constraints. This allows EvoBind to focus designs towards the binding site associated with receptor activation, even in the absence of explicit annotations or structural templates. **b)** Loss distribution and length using the top 10% of designs (n=2500 per length) selected according to equation 1 (Methods). Longer lengths are preferred here, resulting in lower losses, consistent with the length of 31 amino acids of the wildtype (Liraglutide) and that of the agonist in PDB ID 8JIQ used for the comparison here (29 amino acids). Likely, too short peptides do not occupy a sufficient portion of the interface to form a stable interaction. **c)** Predicted confidence (plDDT, higher is better) distribution and length using the top 10% of designs (n=2500 per length) selected according to equation 1. Longer lengths are preferred here, resulting in higher plDDT, consistent with the length of 31 amino acids of the wild type agonist. **d)** Predicted structures of the synthesised top designs of length 15, 18 and 31 (blue, cyan and orange spheres, respectively) in complex with GCGR (green sticks). The predictions are shown in structural superposition with the wild type (WT, yellow) from PDB ID 8JIQ (29 amino acids, designed from GLP1 to be a dual agonist) in stick format. The designs all go down into the binding pocket regardless of length, partially overlapping with the activation surface of the WT binder. **e)** *GCGR agonist response curves*. Agonist-induced cAMP production is measured as relative light units (RLU) in a cell-based assay, plotted against peptide concentration (log M). Data represent means ± standard deviation from three replicates. Designs of length 18 and 31 activate GCGR with EC_50_ values of 542.3 nM and 14.3 μM, respectively, while the 15mer does not elicit any measurable cAMP response. **f)** *GLP1R agonist response curves*. Agonist-induced cAMP production is measured as RLU for GLP1R across peptide concentrations (log M). Data represent means ± standard deviation from three replicates. All three designed peptides induce robust cAMP signalling, comparable to the wild-type agonist Liraglutide (EC_50_=2.4 nM). The length-18 peptide shows the strongest activation (EC_50_=31.9 nM), while the 15mer and 31mer activate GLP1R with EC_50_ values of 11.8 μM and 5.0 μM, respectively.

EvoBind was run across 33 different peptide lengths (8-40 residues), with 5 random seeds per length and 5000 iterations per run, totalling 825000 design steps (Methods). Despite no structural input, EvoBind consistently targeted the extracellular activation surface of GCGR, overlapping the site bound by known peptide agonists (**Figure 1**). This suggests that sequence-to-structure models can internalise functional priors and autonomously identify activation-relevant interfaces in design settings.

Longer peptides showed lower design losses and higher predicted confidence (plDDT [18,20]) scores (**Figure 1b&c**), echoing the length of natural agonists: the GCGR-bound agonist in 8JIQ is 29 amino acids. Yet, cyclic peptides adopt compact topologies, engaging far less receptor surface than linear counterparts, making functional activation especially demanding. We selected the top 10% of sequences according to our loss function (n=82500), filtered for plDDT ≥ 85, yielding 20387 candidates. These were further evaluated with AFM to check for adversarial designs. The five top-scoring sequences were selected for experimental testing.

### Dual Activation of GCGR and GLP1R

GCGR and GLP1R are closely related class B1 GPCRs involved in metabolic regulation, and dual agonists targeting both have shown superior therapeutic effects in obesity and type 2 diabetes [1,21–24]. To explore whether our de novo designed cyclic peptides could achieve dual activity, we evaluated their ability to activate GLP1R, despite having designed them solely against GCGR.

All top-ranked designs were predicted to bind deep within the GCGR functional binding pocket, overlapping the site used by known agonists (**Figure 1d, Methods**). Strikingly, this convergence occurred without any structural templates or binding-site information. The shorter peptides (15 and 18 residues) occupied much smaller regions than native ligands, raising the possibility of generating compact, minimal activation modes through AI-based design.

We synthesised three peptides (lengths 15, 18, and 31) and evaluated their activity in cAMP-based cellular assays against both GCGR and GLP1R (Methods; **Figures 1e&f**). All three activated GLP1R, while the 18mer and 31mer also activated GCGR. The 18mer exhibited the strongest dual activity, with EC_50_ values of 32 nM at GLP1R and 542 nM at GCGR, unprecedented for a de novo designed cyclic peptide agonist. At GLP1R, the 18mer approached the potency of the known agonist Liraglutide [25] derived from the natural GLP1 hormone (EC_50_=2.4 nM), and all peptides induced robust cAMP responses comparable in amplitude to Liraglutide.

The predicted structures of the peptide-GCGR complexes suggest distinct binding modes that, nonetheless, induce receptor conformations matching the active state (RMSD ∼2 Å; **Figure 2a)**. These results demonstrate the feasibility of designing potent, multi-target GPCR agonists with novel activation modes. The fact that the design is blind, i.e. from sequence information alone, suggests its generalisability and introduces cyclic peptides as a promising new modality for GLP1R/GCGR-targeted therapeutics.

**Figure 2.**
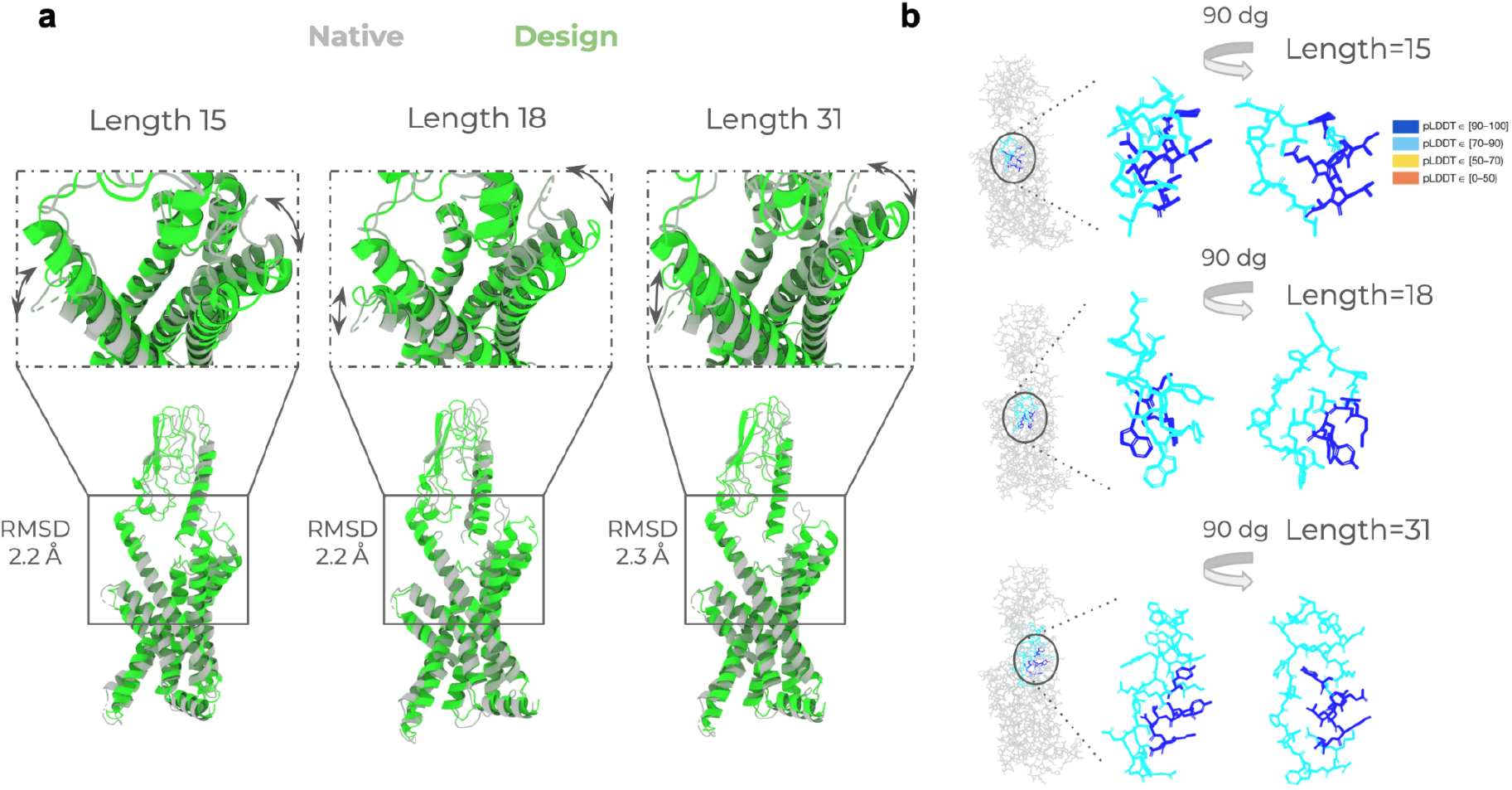
Conformational changes and predicted model confidence in GCGR-peptide complexes. **a)** Overlay of predicted GCGR–peptide complex structures with the active-state GCGR conformation (PDB ID: 8JIQ). Each peptide adopts a distinct binding pose within the orthosteric pocket, inducing receptor conformations closely matching the active state (RMSD ∼2 Å). **b)** Predicted structures coloured by per-residue plDDT scores, representing model confidence. Residues inserted deep into the GCGR binding pocket display the highest confidence levels (plDDT > 90, dark blue), indicating reliable placement at the interface.

**Figure 3.**
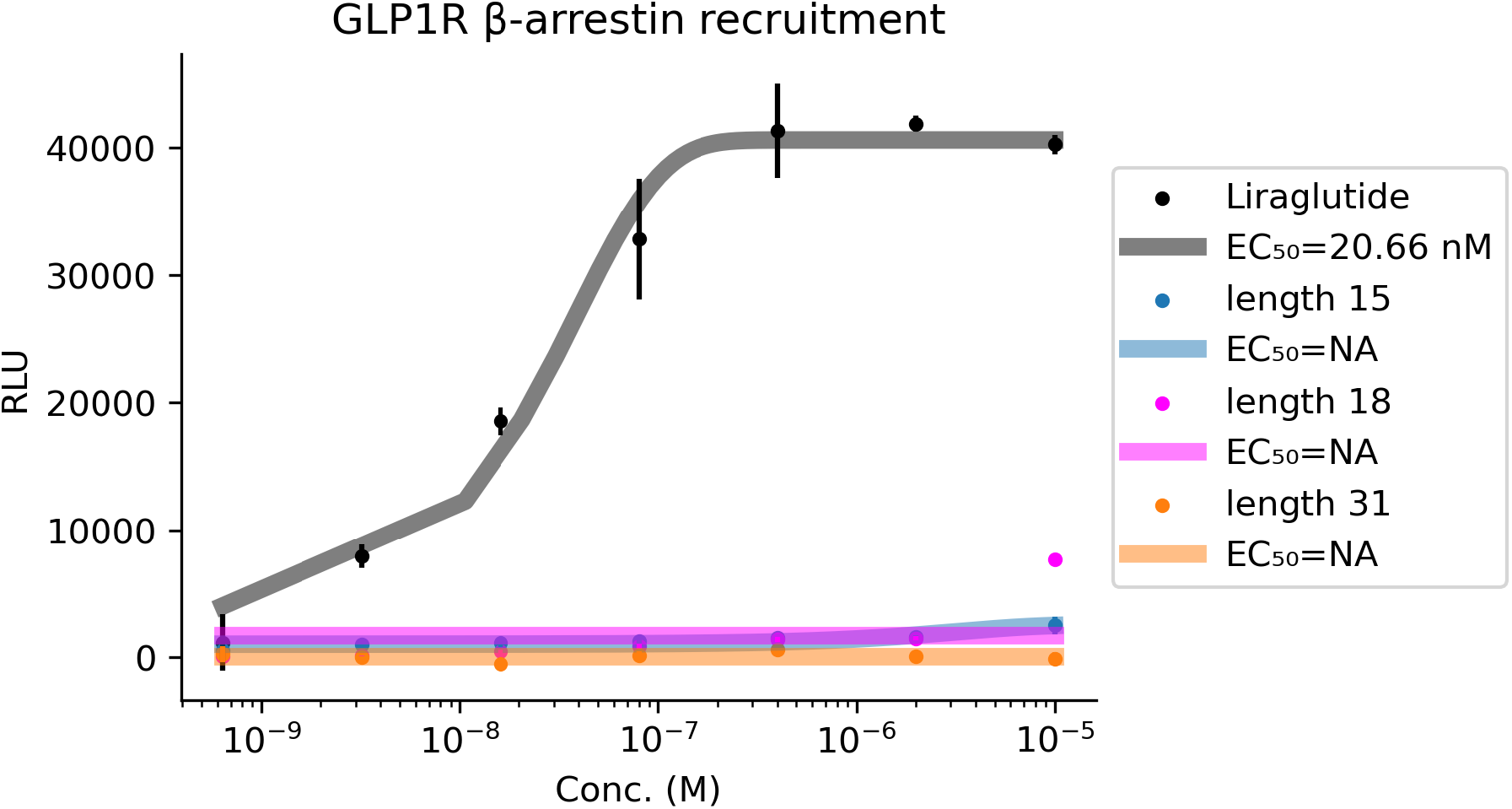
β-arrestin recruitment at GLP1R by designed cyclic peptides compared to Liraglutide. The points mark the averages, and the vertical lines the standard deviation from two replicates, while the fitted lines are used to calculate EC_50_ values. None of the designed peptides elicited substantial β-arrestin recruitment, in contrast to the known agonist Liraglutide, which displayed an EC_50_ of 20.66 nM. The 18-residue peptide, despite strong cAMP activation, showed minimal β-arrestin engagement, highlighting a potential for G protein-biased signalling. Such signalling profiles have been associated with reduced receptor desensitisation and improved glycaemic control in preclinical studies, and have also been observed in recently developed small molecule GLP1R agonists with favourable clinical outcomes.

### Predicted Conformational Modulation Induced by the Designed Cyclic Peptides

The designs of length 18 and 31 exhibit strong GCGR agonist activity, but the 15mer does not, despite all peptides being predicted to bind GCGR. To investigate potential structural explanations, we analysed the predicted GCGR-peptide complexes and associated model confidence (**Figure 2**). All peptides induce conformational changes in GCGR, with overlays against the active-state structure (PDB ID: 8JIQ) shown in **Figure 2a**. Each design adopts a distinct pose within the functional active site, and the cyclic architecture appears to facilitate local rearrangements that open the binding pocket to accommodate diverse conformations. Despite these differences, all peptides induce receptor conformations closely resembling the active GCGR state, with RMSDs around 2 Å.

In **Figure 2b**, peptides are coloured by per-residue plDDT [18], illustrating model confidence at the interface. Residues extending deep into the GCGR binding pocket tend to have high plDDT scores (>90, dark blue), indicating confident placement. However, the number and distribution of such high-confidence residues vary between designs and do not clearly align with functional activity. These results suggest that predicted structural differences and static confidence metrics like plDDT alone are insufficient to explain the observed differences in receptor activation.

The primary distinguishing feature among the designs is peptide length, suggesting that the 15mer may simply be too short to fully engage the GCGR binding pocket. Interestingly, despite lacking measurable GCGR activation, the 15mer elicits GLP1R cAMP signalling comparable to the 31mer, despite not being designed for this target. This discrepancy underscores limitations of current structure prediction methods in capturing subtle determinants of agonist efficacy. Nevertheless, the ability of these de novo cyclic peptides to activate both receptors marks a significant advance toward universal GPCR engagement through sequence-based design.

### GLP1R β-arrestin recruitment

In addition to the cAMP readout for GCGR and GLP1R, we also measured β-arrestin (β-arrestin-2) recruitment for GLP1R. This readout is particularly valuable, as β-arrestin signalling at GLP1R has been implicated in biased agonism, receptor internalisation, and prolonged insulinotropic effects. These downstream effects are not just mechanistically important, but also increasingly therapeutically relevant. Recent strategies in diabetes care explicitly seek to downregulate β-arrestin recruitment at GLP1R to fine-tune efficacy and minimise adverse effects [26]. GLP1R β-arrestin recruitment, therefore, serves as an informative model for assessing the therapeutic potential of novel agonists [27].

We find that none of the designed peptides trigger substantial β-arrestin recruitment at GLP1R compared to the known agonist Liraglutide which reports an EC_50_ value of 20.66 nM. Notably, studies in cellular and animal models suggest that reduced β-arrestin signalling helps to limit receptor desensitisation and downregulation, potentially enhancing anti-hyperglycaemic efficacy. Against this backdrop, the ability of our peptides, particularly the 18mer, to strongly activate the cAMP pathway while not recruiting β-arrestin highlights their therapeutic potential and mechanistic selectivity [26]. Similar effects, strong cAMP activation and lacking β-arrestin recruitment, have also been observed preclinically in recent small-molecule GLP1R agonists with a subsequent positive outcome in humans [28].

## Discussion

This work demonstrates the first successful case, to our knowledge, of *blind de novo design* of a cyclic peptide agonist for a G protein-coupled receptor (GPCR), using only the target receptor sequence as input. This outcome highlights the power of EvoBind [15] and the broader potential of *in silico* directed evolution and protein design for therapeutic discovery.

While AI-based drug design is still in its infancy and often met with warranted scepticism [29,30], especially when no structural information or epitope guidance is provided, our results provide evidence that functional receptor activation can emerge from blind sequence-based design. Importantly, the task of designing not just binders but protein activating *agonists* is significantly more demanding: it requires the peptide to induce the specific conformational rearrangements in the receptor associated with intracellular signalling. This is particularly challenging for the class of GPCRs that GCGR and GLP1R belong to, whose activation relies on highly specific interactions with the transmembrane core [31,32].

We show that several designed cyclic peptides activate both GCGR and GLP1R, with the most potent 18-residue design achieving nanomolar EC_50_ values, comparable to Liraglutide, a clinically used GLP1 analogue derived from the endogenous hormone. Remarkably, our peptides achieve this potency despite adopting predicted binding modes that are necessarily distinct from those of natural ligands or existing drugs. This is supported by their divergent sequences and cyclic topology, which are incompatible with the known orthosteric binding sites. We underline that sequence-based frameworks are essential if novel binding modes are to be discovered, a necessity for accommodating cyclic peptides. Although the design process targeted only GCGR, and the broader applicability of this approach remains to be established, the observed cross-activation of GLP1R suggests that purely sequence-based design can enable the generation of multi-target or broadly active agonists, particularly within families of structurally related receptors.

Interestingly, the designed peptides potently activate GLP1R in cAMP assays but do not induce β-arrestin recruitment, unlike Liraglutide, which robustly engages both pathways. This selective signalling profile is of particular interest, as reduced β-arrestin engagement has been linked to decreased receptor desensitisation and downregulation, offering a potential path to improved long-term efficacy. The 18-residue peptide, in particular, demonstrates strong cAMP activation with minimal β-arrestin signalling, closely resembling the profile of recently developed G protein-biased small molecule agonists that have shown favourable outcomes in humans [28]. This suggests that EvoBind’s sequence-based design process may not only uncover novel agonists but also favour signalling selectivity with therapeutic relevance.

Our previous work has demonstrated that EvoBind can generate both linear and cyclic peptide binders across diverse targets and lengths, achieving nanomolar affinity in a single generative step [15,16]. Recent extensions incorporating noncanonical amino acids [33] promise to expand its design capabilities even further. In this study, the successful creation of de novo receptor agonists with distinct activation profiles reinforces the idea that purely sequence-based protein structure prediction and EvoBind’s directed evolution approach can uncover functional interaction surfaces, not just static binders. Given the inherent challenges of GPCR activation, these results provide a compelling proof-of-concept for target-agnostic cyclic peptide design as a powerful and general strategy for therapeutic discovery.

## Methods

### Cyclic peptide design

For designing cyclic peptide binders, we extracted the sequence from the entire GLP1R (PDB: https://www.rcsb.org/structure/8jiq, chain C) and input it to EvoBind (version 2) [15]. EvoBind explores a relaxed sequence-structure space by mutating one residue at a time to minimise the following loss function:

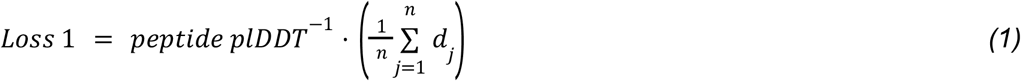

The peptide pLDDT represents the average predicted LDDT score across all peptide residues. For each atom j in the peptide, dj denotes the shortest distance to any atom in the receptor, and nn is the total number of atoms in the peptide.

To facilitate the design of cyclic peptides, we incorporated a cyclic offset into the design protocol. This approach, inspired by previous work [14], modifies the input to the structure prediction networks to treat the peptide sequence as a continuous cycle rather than a linear chain. This cyclic offset was applied for both AlphaFold2 [18,34] and AlphaFold-multimer [19] networks (below).

The peptide is represented by a single sequence, initiated using a Gumbel distribution over all 20 standard amino acids. The GLP1R is represented by a multiple sequence alignment (MSA) constructed from searching uniclust30_2018_08 [35] with HHblits version 3.1.0 [36] using the following command:

`hhblits -E 0.001 -all -oa3m -n 2`

For folding and optimisation, EvoBind utilises the single-chain AlphaFold2 folding pipeline with model_1, one ensemble, and 8 recycles. The optimisation process ran for 5000 iterations and was repeated five times from random starting points for exploration. We explored a diverse set of lengths, from 8-40 residues, resulting in a total of 5000⋅5⋅33=825000 predictions. See **Figures 4 and 5** for the optimisation curves.

**Figure 4.**
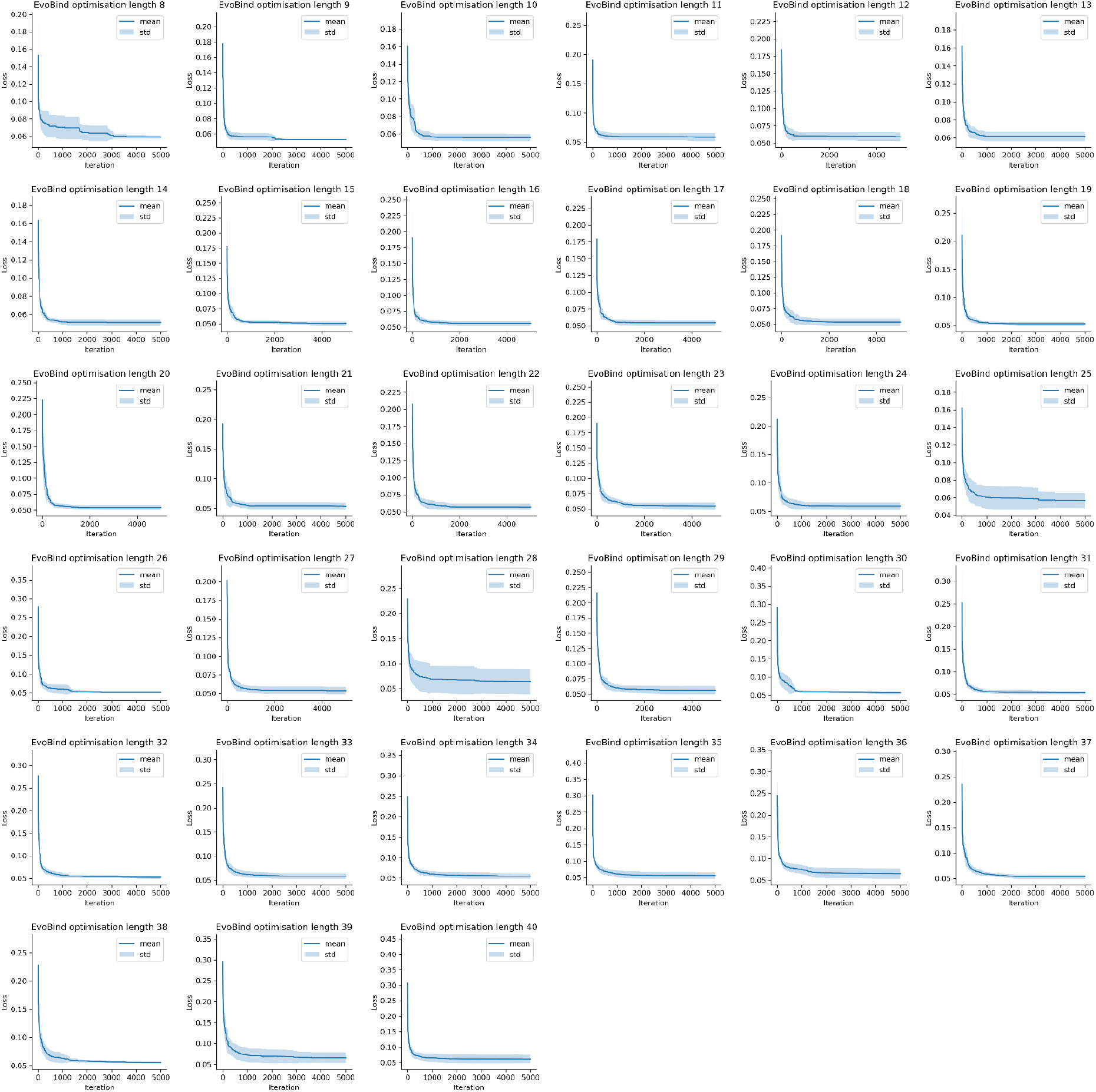
Loss optimisation curves across design trajectories. EvoBind optimisation was run for 5000 iterations per trajectory, repeated five times from independent random initialisations to enhance exploration. Designs spanned 33 peptide lengths ranging from 8 to 40 residues, yielding a total of 825000 structure predictions (5,000 ⋅ 5 ⋅ 33). The curves show the loss trajectories over iterations for all lengths, illustrating consistent convergence and effective optimisation across runs. The means are marked by solid lines, and the standard deviations are shaded. The best loss at each step is shown. Most losses converge within 1000 iterations, indicating that the 5000 iterations used here may be reduced substantially.

**Figure 5.**
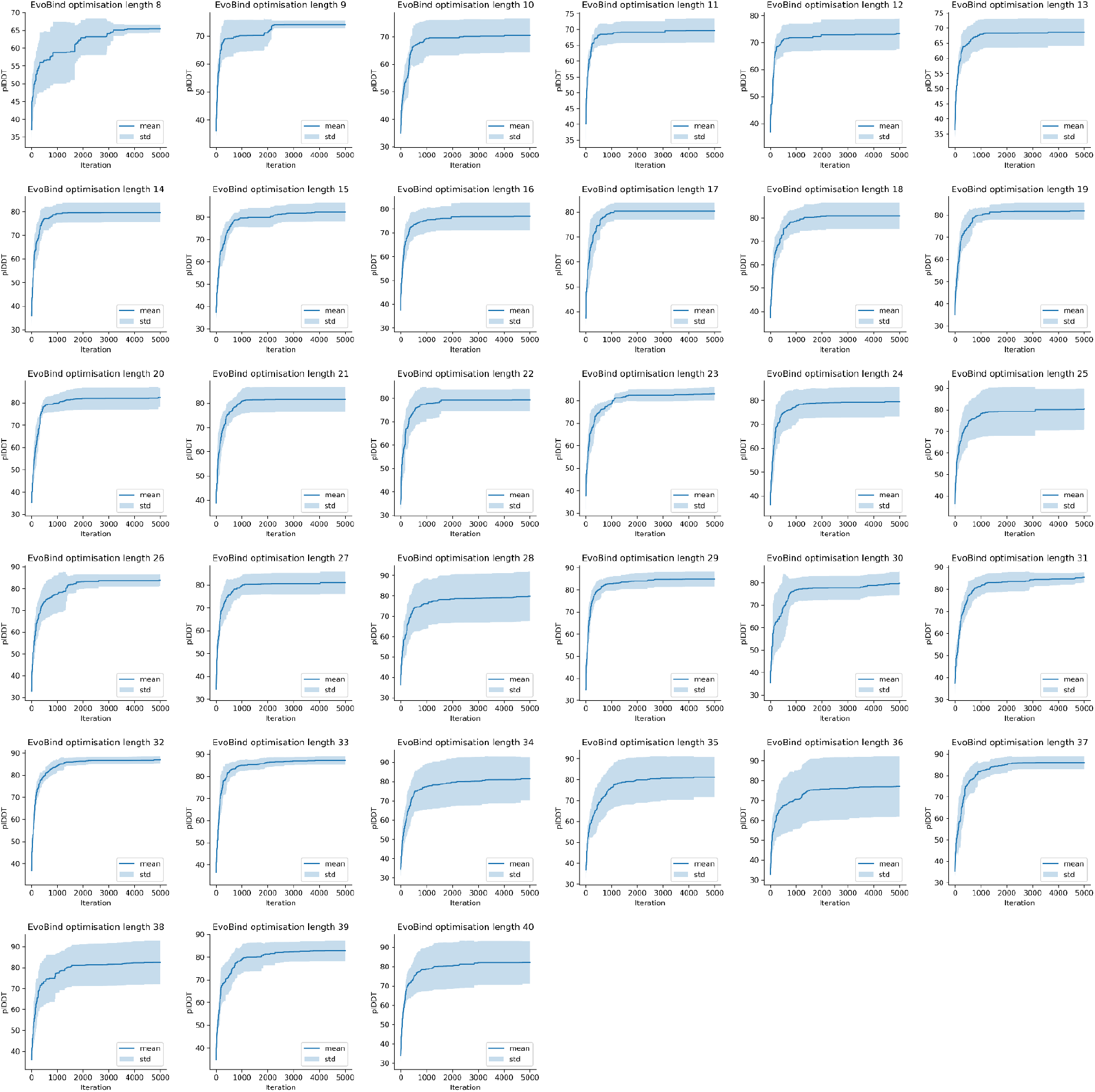
plDDT optimisation curves across design trajectories. EvoBind optimisation was run for 5000 iterations per trajectory, repeated five times from independent random initialisations to enhance exploration. Designs spanned 33 peptide lengths ranging from 8 to 40 residues, yielding a total of 825000 structure predictions (5,000 ⋅ 5 ⋅ 33). The curves show the plDDT trajectories over iterations for all lengths, illustrating consistent convergence and effective optimisation across runs. The means are marked by solid lines, and the standard deviations are shaded. The best plDDT at each step is shown. Most plDDTs converge within 1000 iterations, indicating that the 5000 iterations used here may be reduced substantially.

### Selection of designs for experimental validation

From the generated designs, we selected the top 10% of the sequences based on equation 1 (n=82500) and from these, all designs with an average peptide plDDT≥85 (n=20387). A secondary evaluation using AlphaFold-Multimer was then conducted to avoid adversarial designs, which has previously been shown to improve the success rate [13] substantially. The AFM configuration uses version v2.1.0 (params_model_1_multimer_v2) with 20 recycles and early stopping if the predicted confidence score doesn’t improve. Dropout is applied everywhere except in the structural model. The similarity between the structures predicted by EvoBind2 and AlphaFold-Multimer (AFM), along with the confidence of the AFM prediction (measured by the plDDT), is used to calculate a second loss:

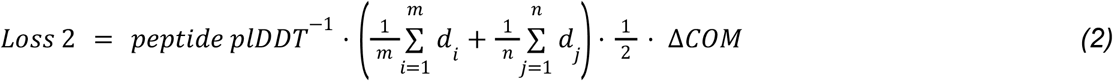

Here, the peptide plDDT refers to the average plDDT across the peptide; didi is the shortest distance between any atom in the receptor target atoms (specifically, Cβ atoms within 8 Å of the peptide, as predicted by EvoBind2) and any atom in the peptide; dj is the shortest distance between any atom in the peptide and any atom in the receptor target atoms; and ΔCOM denotes the distance between the Cα centres of mass of the predicted peptides from the design and validation procedures, respectively.

We select the top designs (lowest combined loss) from the best five unique lengths for GCGR that have plDDT≥85 (**Table 1**). We excluded the solubility filter previously used in EvoBind as these are based on the fraction of charged and hydrophobic residues being above/below 25%, however the agonist in GCGR/GLP1R (PDB ID 8JIQ/8JIS) has 24% charged and 31% hydrophobic residues (HSQGTFTSDYSKYLDEQAAKEFIAWLMNT). The longest lengths (37 and 38) could not be easily synthesised, therefore, only lengths 15, 18 and 31 were evaluated experimentally.

**Table 1.** Selection for GCGR. The table columns represent key metrics from peptide design and characterisation. **if_dist_peptide** indicates the interface distance between peptide and receptor in predicted complexes, reflecting binding proximity. **plddt_evo** and **plddt_afm** are confidence scores from EvoBind and AlphaFold-Multimer (AFM) structural predictions, respectively, while **loss_evo** (equation 1) and **loss_afm** (equation 2) quantify the corresponding model losses, with lower values indicating better predicted interactions. **sequence** lists the amino acid sequences of designed cyclic peptides. **COM** measures the center of mass distance between EvoBind and AFM peptide predictions, with values below 1 Å signifying high structural similarity. **combined_loss** is the average of both EvoBind and AFM losses to assess overall design quality. Finally, **Purity** reflects the experimentally measured purity of synthesized peptides, expressed as a percentage.

| run | iteration | length | if_dist_peptide | pLDDT_evo | loss_evo | sequence | loss_afm | pLDDT_afm | COM | combined_loss | Purity | GCGR EC50 | GLP1R EC50 |
| --- | --- | --- | --- | --- | --- | --- | --- | --- | --- | --- | --- | --- | --- |
| 1 | 2801 | 15 | 4,3 | 85 | 0,051 | YIIQTLALDPDPNIP | 0,051 | 90 | 0,4 | 0,102 | 96.9% | NA | 11.77 μM |
| 4 | 2047 | 18 | 4,2 | 86 | 0,048 | LCWYGYNDPQAP<br>AMVNWY | 0,073 | 86 | 0,6 | 0,122 | 95.8% | 542.34 nM | 31.86 nM |
| 5 | 2010 | 31 | 4,4 | 86 | 0,051 | CQAKYQEYTDKIN<br>MQGWNVPELYCN<br>DPHSWN | 0,010 | 85 | 0,1 | 0,062 | 90.2% | 14.33 μM | 5.02 μM |
| 2 | 4763 | 37 | 4,6 | 86 | 0,053 | REAGTCPLWEADA<br>IQCIASYAIVKNSPL<br>GDSMKAGIW | 0,031 | 86 | 0,3 | 0,084 | NA | NA | NA |
| 1 | 2645 | 38 | 4,7 | 86 | 0,054 | EGPNWRNAMAGK<br>QKWVIDSQGYVYT<br>VNMTYEAVHAISA | 0,062 | 86 | 0,7 | 0,116 | NA | NA | NA |

To validate the structural consistency of our designs, we compared EvoBind (AF2-based) predictions with independent AlphaFold-Multimer (AFM) models of the same peptide-receptor complexes. The predicted binding poses showed a high degree of overlap, with the centres of mass (COM) for the peptides aligning within less than 1 Å (Table 1, **Figure 6**). This close correspondence confirms that both methods agree on similar binding modes, reinforcing the reliability and accuracy of EvoBind’s de novo design through this adversarial evaluation.

**Figure 6.**
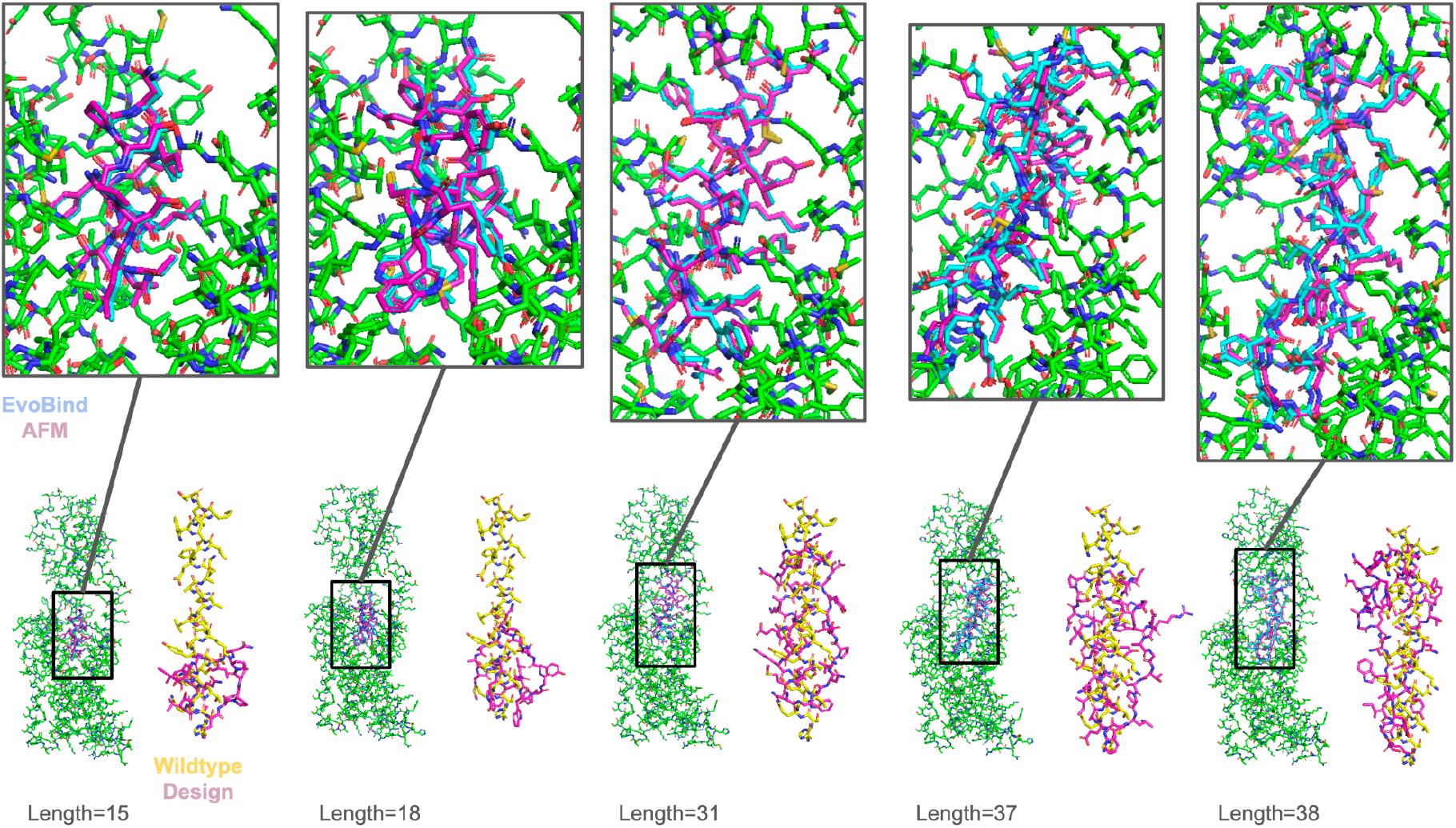
Comparison of top-ranked peptide-GCGR complex predictions from EvoBind using AlphaFold2 (AF2) and AlphaFold-Multimer (AFM). The binding poses of all top designs are highly similar between the two prediction methods, with peptide centers of mass (COM) differing by less than 1 Å. This strong concordance highlights the robustness and reproducibility of the predicted binding modes across independent structure prediction frameworks.

When comparing the sequences of the three successfully synthesised peptides to that of the dual agonist from 8JIQ, there is almost no similarity in terms of consecutive sequence similarity, underlining the novelty of our cyclic designs **(Figure 7)**.

**Figure 7.**
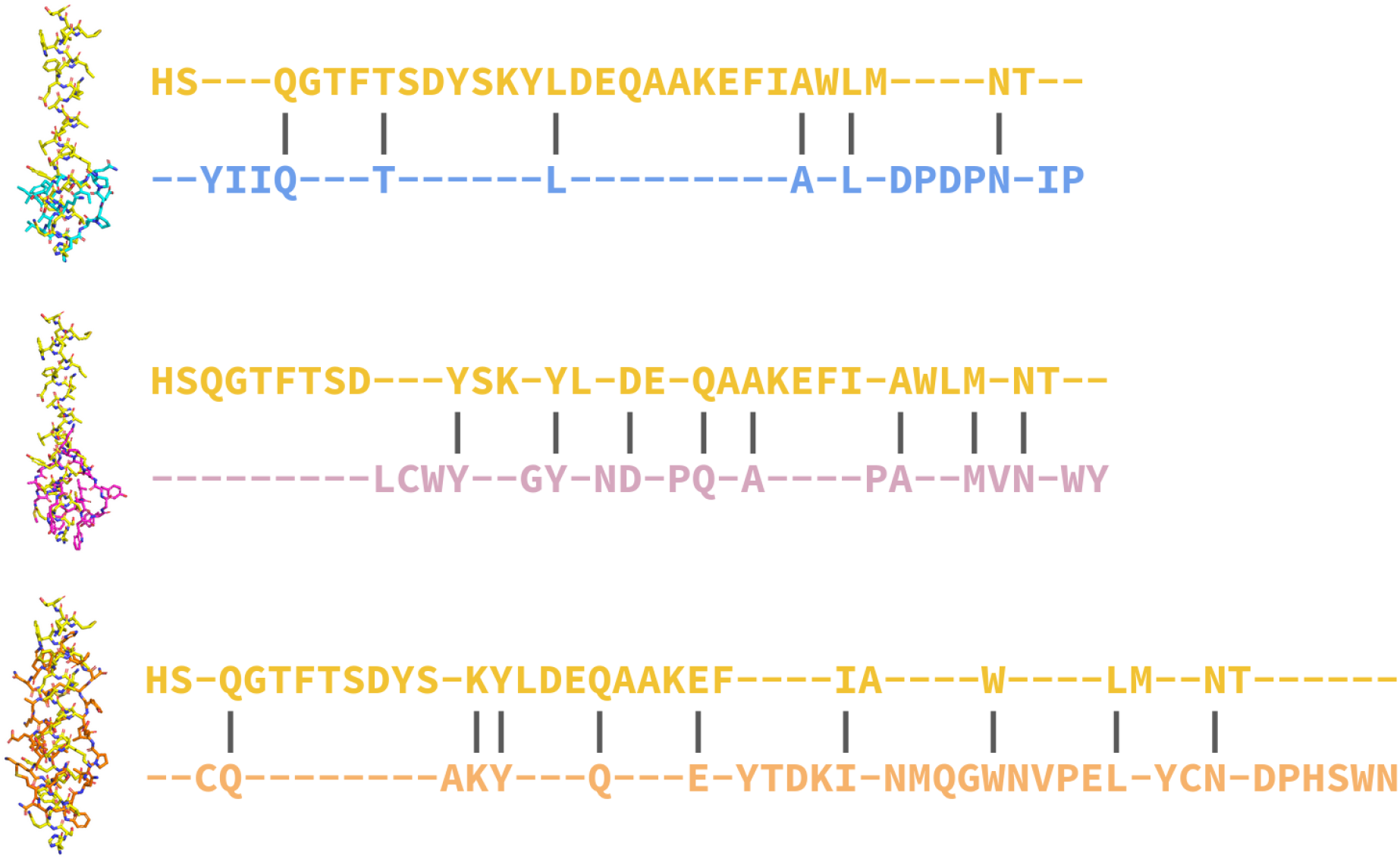
Global sequence alignments comparing the three top designs of lengths 15, 18 and 31 to the known binder in PDB ID 8JIQ. There is little sequence similarity with the known binder, indicating the novelty of the designs. The binder is an analog to the natural GLP-1 peptide hormone (HAEGTFTSDVSSYLEGQAAKEFIAWLVKGRG).

### GCGR target sequence used for design

#### GCGR: https://www.rcsb.org/structure/8jiq

>8jiq_R QVMDFLFEKWKLYGDQCHHNLSLLPPPTELVCNRTFDKYSCWPDTPANTTANISCPWYLPWHHKVQHRFVFKRCGPDGQWVRGPRGQPWRDASQCQMDGEEIEVQKEVAKMYSSFQVMYTVGYSLSLGALLLALAILGGLSKLHCTRNAIHANLFASFVLKASSVLVIDGLLRTRYSQKIGDDTWLSDGAVAGCRVAAVFMQYGIVANYCWLLVEGLYLHNLLGLATLPERSFFSLYLGIGWGAPMLFVVPWAVVKCLFENVQCWTSNDNMGFWWILRFPVFLAILINFFIFVRIVQLLVAKLRARQMHYKFRLAKSTLTLIPLLGVHEVVFAFVGTLRSAKLFFDLFLSSFQGLLVAVLYCFLNKEVQS ELRRRWHRW

### Peptide synthesis and preparation

Lyophilised powders were synthesised and purified by GenScript. The quality of the peptide (purity above 90%) was verified by high-performance liquid chromatography and mass spectroscopy (see the Supplementary files).

Cyclic peptides length 15 and length 31 were dissolved in Milli-Q water, while the cyclic peptide length 18 was dissolved in DMSO. All peptides were diluted to a concentration range of 10 µM to 0.001 nM for the β-arrestin assay and 30 µM to 1.8 nM for the cAMP assay, based on net peptide content. The final DMSO concentration in the cell assays was kept consistently at a maximum of 0.1% to avoid adverse effects and cell death.

### Liraglutide

We ordered Liraglutide (Cat. No.: HY-P0014) from MedChemExpress. This contains 1 mg Liraglutide dissolved in 1 mL sterile filtered 10mM PBS, pH 7.4.

### Net Peptide Content Determination

The peptide was lyophilised with TFA salt and water. To analyse the actual peptide content, we diluted the lyophilised powder to 1 mg/mL for length 15 and length 31 in Milli-Q water and 11 mg/mL for length 18 in DMSO (due to lower solubility reported by GenScript, see the Supplementary files), and used UV spectroscopy with Nanodrop. The designed sequences all contain aromatic amino acids that absorb light at 280 nM (see below).

The absorbance is related to the concentration (c) and path length (l=10 mm) in the following way:

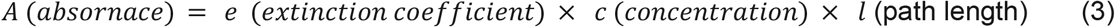

In detail, this is how the extinction coefficients in Table 2 were calculated for each sequence:

**Table 2.**
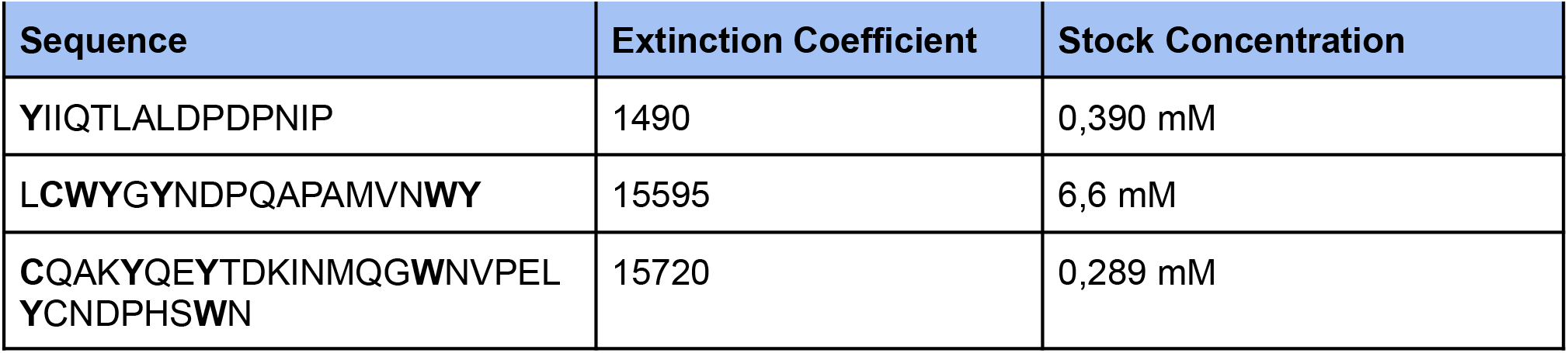
Extinction coefficients and calculated stock concentrations for each peptide based on the measured absorbance.

**Y**IIQTLALDPDPNIP

The extinction coefficients are calculated accordingly:

1xTYR (Y, e = 1490)

The extinction coefficient: 1490

We measured the absorbance value is 0.584 at 280 nm. The absorbance is related to the concentration (c) and path length (l=10 mm) in the following way:

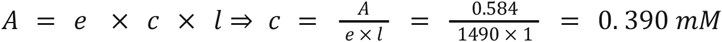

L**CWY**G**Y**NDPQAPAMVN**WY**

**1xCYS (C, e = 125) + 2xTRP (W, e = 5500) +** 3xTYR (Y, e = 1490)

The extinction coefficient: 1 × 125 + 2 × 5500 + 3 × 1490 = 15595

We measured the absorbance value is 103.9 at 280 nm. The absorbance is related to the concentration (c) and path length (l=10 mm) in the following way:

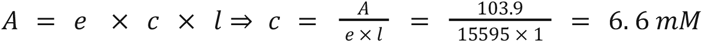

**C**QAK**Y**QE**Y**TDKINMQG**W**NVPEL**Y**CNDPHS**W**N

**2xCYS (C, e = 125) + 2xTRP (W, e = 5500) +** 3xTYR (Y, e = 1490)

The extinction coefficient: 2 × 125 + 2 × 5500 + 3 × 1490 = 15720

We measured the absorbance value is 4.542 at 280 nm. The absorbance is related to the concentration (c) and path length (l=10 mm) in the following way:

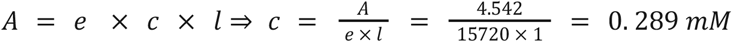

### Cyclic adenosine monophosphate accumulation assay

To quantify cellular cAMP levels, cAMP Hunter eXpress GPCR Kits were used (Qiagen, 95-0062E2CP2L and 95-0042E2CP2L). CHO K1 GCGR/GLP1R cells were plated onto 384-well plates (BD Falcon, 353962) and incubated overnight. Cells were then stimulated with controls (Liraglutide) and designed cyclic peptides for 1 hour at 37°C and further processed according to the manufacturer’s protocol. Briefly, following agonist incubation, 15 μL of cAMP Antibody Reagent was added to all wells of the assay plate. A cAMP Working Detection Solution was prepared in a separate 15 mL polypropylene tube by mixing 19 parts cAMP Lysis Buffer, 5 parts Substrate Reagent 1, 1 part Substrate Reagent 2, and 25 parts cAMP Solution D. Subsequently, 60 μL of the cAMP Working Detection Solution was added to all wells without pipetting up and down or vortexing the plates. The assay plate was incubated for 1 hour at room temperature in the dark to allow the immunocompetition reaction to occur. Next, 60 μL of cAMP Solution A was added to all wells without pipetting up and down or vortexing, followed by a 3-hour incubation at room temperature in the dark. Luminescence was measured using a standard luminescence plate reader with a read time of 1 second per well for the photomultiplier tube.

### Curve fitting and EC_50_ calculation

Dose-response curves were fitted to the RLU measurements using a four-parameter sigmoidal function of the form:

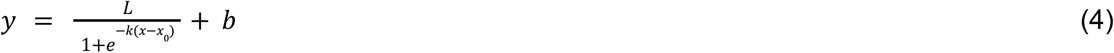

Where *L* is the response range, *k* is the slope, *x*_0_ is the inflexion point, and *b* is the baseline. Curves were adjusted using negative controls (measured RLU without peptide addition), and EC_50_ values reported here were calculated from these fits.

### Plasmid preparation

GLP1R-Tango and GCGR-Tango were gifts from Bryan Roth (Addgene plasmid # 66295 ; http://n2t.net/addgene:66295 ; RRID:Addgene_66295; (Addgene plasmid # 66291 ; http://n2t.net/addgene:66291 ; RRID:Addgene_66291) [37]. To purify this plasmid, bacteria (DH5α) containing the plasmid were cultured overnight at 37°C in LB medium supplemented with 100 μg/mL of antibiotic. The plasmid DNA was then purified using the PureLink™ Quick Plasmid Miniprep Kit according to the manufacturer’s instructions. The concentration of the purified plasmid DNA was measured using a NanoDrop spectrophotometer.

### Presto-Tango assay

HTLA cells (HEK293 cell line stably expressing a tTA-dependent luciferase reporter and a β-arrestin-TEV protease fusion gene) were a gift from Bryan Roth, were cultured in DMEM supplemented with 10% foetal bovine serum (FBS), 100 μg/mL of penicillin-streptomycin, 2 μg/mL of puromycin and 100 μg/mL of hygromycin at 37 °C in a humidified atmosphere containing 5% CO2.

For transfection, the HTLA cells were plated in 100 mm dishes at 6×10^6^ cells/dish in 10 ml culture medium. The following day, 20 µg of GLP1 plasmid DNA was transfected into cells using the Calcium Phosphate Transfection kit (Invitrogen #K2780-01). Transfected cells were transferred at 15,000 to 20,000 cells per well in 40 μl of medium into poly-D-lysine coated and rinsed 384-well white clear-bottom as the cell culture plates (Greiner Bio-one). Stock peptides were diluted into various concentrations with an assay buffer (HBSS+20 mM HEPES), and dispensed with SELMA384 into the cell culture plate. The 55 µl/well medium and peptide solutions were removed from the wells with SELMA 384. Bright Glo, diluted 1:10 with assay buffer, was added with Multidrop Combi (low speed), 20 µl/well. Plates were incubated 15-20 min at room temperature in the dark followed by luminescence readings in Hidex Sense.

## Availability

Data: https://zenodo.org/uploads/13933365 Code: https://github.com/patrickbryant1/EvoBind

## Contributions

PB designed the study and wrote the initial draft of the manuscript. QL performed binder design and conducted β-arrestin wet lab assays. EW and TH provided additional experimental support. All authors contributed to data analysis and approved the final manuscript.

## Acknowledgements

We thank Bartlomiej Porebski at the Chemical Biology Consortium Sweden for performing the cAMP assays.

